# Fully-sensitive Seed Finding in Sequence Graphs Using a Hybrid Index

**DOI:** 10.1101/587717

**Authors:** Ali Ghaffaari, Tobias Marschall

## Abstract

**Motivation:** Sequence graphs are versatile data structures that are, for instance, able to represent the genetic variation found in a population and to facilitate genome assembly. Read mapping to sequence graphs constitutes an important step for many applications and is usually done by first finding exact seed matches, which are then extended by alignment. Existing methods for finding seed hits prune the graph in complex regions, leading to a loss of information especially in highly polymorphic regions of the genome. While such complex graph structures can indeed lead to a combinatorial explosion of possible alleles, the query set of reads from a diploid individual realizes only two alleles per locus—a property that is not exploited by extant methods.

**Results:** We present the ***P****an-genome* ***S****eed* ***I****ndex (PSI)*, a fully-sensitive hybrid method for seed finding, which takes full advantage of this property by combining an index over selected paths in the graph with an index over the query reads. This enables PSI to find all seeds while eliminating the need to prune the graph. We demonstrate its performance with different parameter settings on both simulated data and on a whole human genome graph constructed from variants in the 1000 Genome Project data set. On this graph, PSI outperforms GCSA2 in terms of index size, query time, and sensitivity.

**Availability:** The C++ implementation is publicly available at: https://github.com/cartoonist/psi.

## 1 Introduction

The reference genome of a species is intended to be the representative genome of its population. The “linear” reference genomes in use today, at best, reflect a consensus genome of all individuals, but do not capture small variants and structural diversity of a population (Church et al., 2015). When mapping reads to such references, this leads to a *reference bias*: reads supporting the reference allele have a higher chance of being aligned compared to reads supporting an alternative allele (Paten et al., 2017; Garrison et al., 2018; Rakocevic et al., 2019). This limitation hampers the performance of downstream analyses such as variant calling. In particular, short reads coming from highly divergent regions, such as the human leukocyte antigen (HLA) genes, often remain unmapped or misaligned (Dilthey et al., 2015).

At the same time, advances in high-throughput sequencing technologies have enabled gathering extensive catalogues of genetic variation, for instance by the 1000 Genomes Project (1KGP) (Auton et al., 2015). With the advent of long read technologies, the *de novo* assembly of individual human genomes has now become feasible, which additionally uncovers substantial amounts of structural variation missed in short-read based studies (Chaisson et al., 2015, 2017; Audano et al., 2019). Importantly, such assembly-based approaches are able to resolve the full sequences of alternative alleles. Translating this growing knowledge about genomic diversity in humans into improved analysis pipelines for (re-)sequencing data constitutes a pressing challenge in bioinformatics.

Consequently, there is a growing interest in data structures capable of representing a species’ *pan-genome*, that is, to encode a comprehensive amount of sequence found in the genomes of a population (Computational Pan-Genomics Consortium, 2018). Pangenomes can be represented in different ways that come with varying computational advantages and limitations. One simple approach consists in augmenting the reference genome with alternative alleles for important loci, a strategy that is implemented (to a limited extend) in the current version of the human reference genome GRCh38 (Church et al., 2015). Graph-based representations, in contrast, can express polymorphisms more flexibly and succinctly, but introduce substantial computational challenges (Paten et al., 2017; Computational Pan-Genomics Consortium, 2018). Initial studies have demonstrated clear benefits, including a reduced reference bias (Garrison et al., 2018; Rakocevic et al., 2019), enhanced variant calling (Eggertsson et al., 2017), as well as improved allele inference of difficult loci such as the HLA genes (Dilthey et al., 2015, 2016).

Despite these successes, considerable algorithmic challenges remain. In particular, we are not aware of any read alignment tool able to map reads to (arbitrary) graphs at speeds comparable to tools for mapping reads to linear sequences. Most read aligners, both for mapping to linear sequences and for mapping to graphs, rely on a *seed-and-extend* approach (Li and Homer, 2010; Reinert et al., 2015). That is, they first find short (exact or approximate) matches, called *seed hits*, and subsequently *extend* these seed hits to obtain alignments. The seed finding step can be fundamentally more challenging on graphs than on sequences, because complex regions in the graph can give rise to a combinatorial explosion in the number of possible paths. Notably, the process of *aligning* reads to graphs is not disturbed by this, and efficient algorithms for aligning sequences to graphs exist (Myers and Miller, 1989; Navarro, 2000; Rautiainen et al., 2019). In this paper, we therefore focus on the seed finding step with a particular focus on handling variant-dense regions in the input graph.

### 1.1 Related Work

Collections of similar sequences can be indexed using Burrows-Wheeler-Transform (BWT) based techniques (Mäkinen et al., 2010), which exploit similarities between the sequences in order to save space. We refer the reader to the review by Gagie and Puglisi (2015) for further discussion of related techniques for indexing collections of sequences and focus on specific techniques to index sequence-labeled graphs in the following.

Most existing indexing schemes for sequence graphs attempt to index *k*-mers in the graph, and they can broadly be categorized as being either hashing-based or BWT-based. The first hashing-based approach was introduced by Schneeberger et al. (2009), and several related approaches based on hashing *k*-mers have been put forward since then (Danek et al., 2014; Limasset et al., 2016; Eggertsson et al., 2017; Petrov et al., 2018).

Instead of hashing methods, *de Bruijn graphs* can be used as a basis for indexing *k*-mers occurring in sequence graphs. The *XBW transform* (Ferragina et al., 2009), which is an extension of the FM index (Ferragina and Manzini, 2005) to labeled trees, has inspired approaches like *Succinct de Bruijn graphs* by Bowe et al. (2012), kFM-index by Rødland (2013), and GCSA by Sirén et al. (2014). Later, GCSA2 (Sirén, 2017) was introduced to improve the original GCSA by employing the ideas of succinct de Bruijn graphs. This modiffication relaxed the constraints on cycles in the graph while it imposed an upper-bound limit on the length of query searches.

The key limitation of all above approaches is the combinatorial explosion of the *k*-mer space as more variants are added to the sequence graph. Thus, the index size can grow exponentially which, consequently, increases the memory footprint and run-time of the read alignment. In order to handle human genomes, these methods therefore need to prune the input graph, which can potentially lead to breaking haplotype paths, or to removing known variants from the sequence graph.

### 1.2 Contributions

In this article, we propose the first scalable, fully-sensitive method for finding seeds in a node-labeled directed graph. We call our approach PSI, which is short for ***P****an-genome* ***S****eed* ***I****ndex*. PSI is a hybrid approach that utilizes the indexes of both reference graph and query reads. We leverage the idea that the *k*-mer space in the read library is much more limited than that in the graph. In particular, it is independent of the number of variants in the graph.

In a preprocessing phase, we construct a collection of paths through the graph and index them using a conventional FM index. Our method for selecting these paths is designed to cover as many *k*-mers present in the graph as possible. Our evaluation shows that this path index alone outperforms GCSA2—a highly optimized indexing method proposed by (Sirén, 2017)—in terms of index size, query time, and sensitivity when indexing all SNVs with allele frequency above 1% found in the 1000 Genomes Project.

Still, our path index does not reach full sensitivity; that is, it misses *k*-mers in variant-dense regions of the graph. We refer to such loci in the graph where the path index misses *k*-mers as *uncovered loci*. To rescue missed *k*-mers at uncovered loci, we index a *chunk* (=subset) of input reads at a time. We then *traverse* the graph, starting from all uncovered loci, and the read index in parallel. The full workflow is illustrated in Figure 1. This approach turns out to be efficient in practice for multiple reasons: (i) by traversing read index and graph simultaneously, *k*-mers that are not represented in the read set are avoided, circumventing extra *k*-mers present in the graph; (ii) even when including all variants from the 1000 Genomes Project, the number of these uncovered loci remains manageable; and (iii) the size of the chunks is a tuning parameter that can be adjusted such that the read index fits into the processor cache, which makes traversing it very fast. As a result, our hybrid indexing strategy reaches full sensitivity at a moderate overhead compared to the path-only index, and is (to our knowledge) the first scalable technique providing full sensitivity for large, variant-dense graphs.

**Figure 1.**
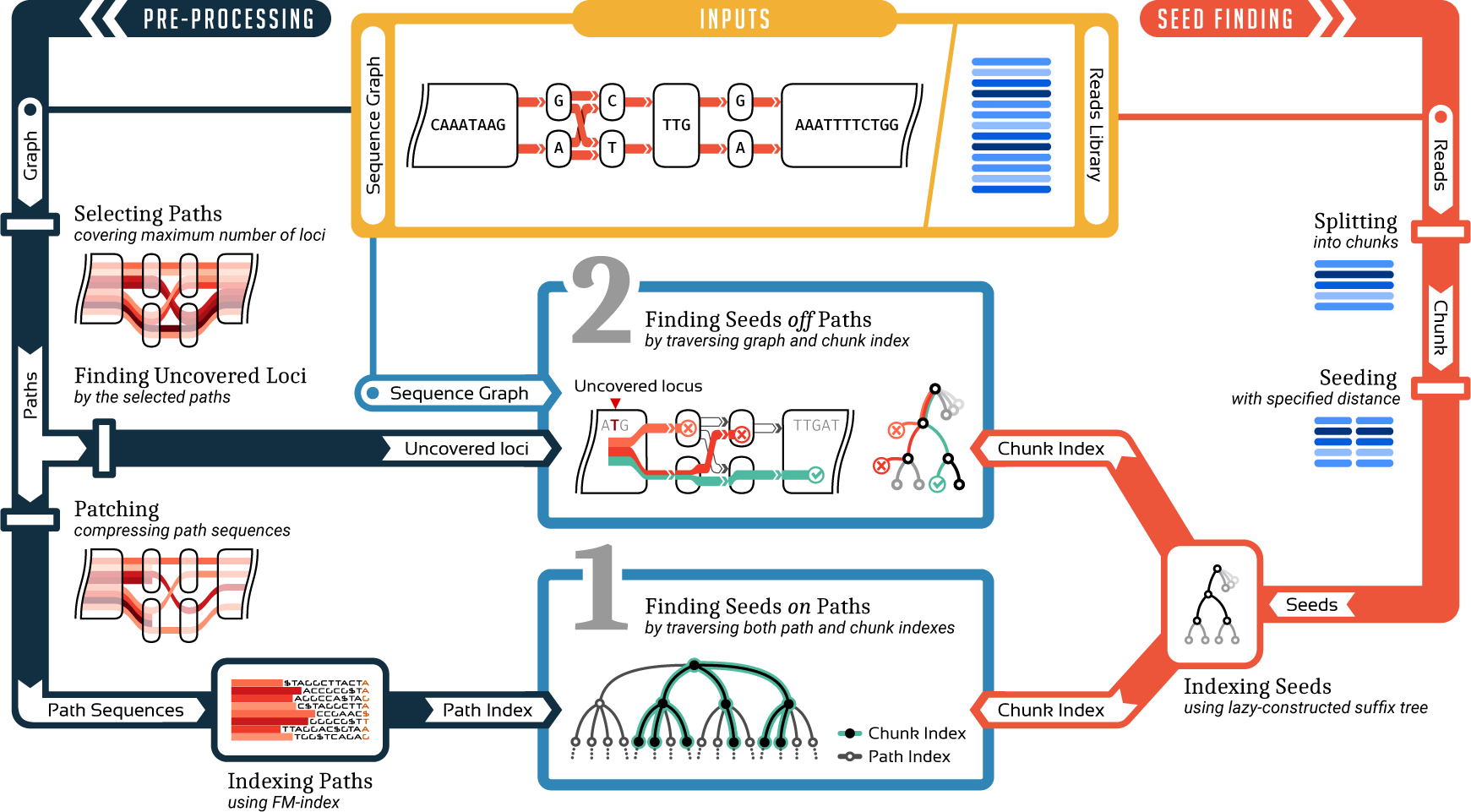
Conceptual overview of the PSI approach. A sequence graph and a set of reads are provided as inputs (yellow). The graph is preprocessed to create a path index (left / dark blue). The read set is split into chunks and seeds are indexed (right / red). Seed finding proceeds in two stages (middle / light blue).

**Figure 2.**
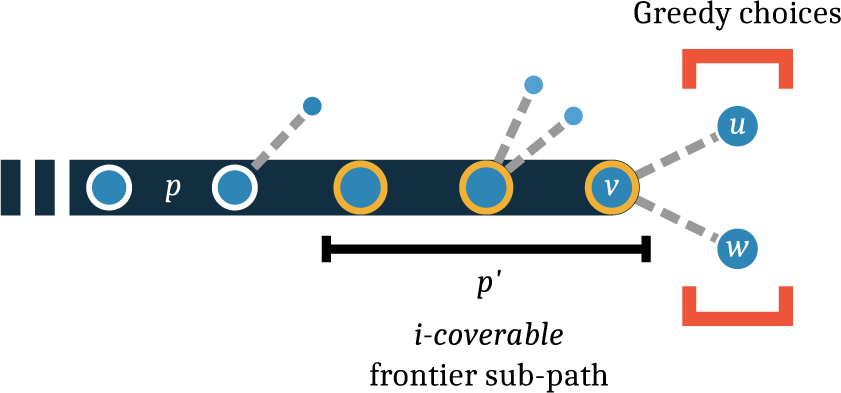
Illustration of path selection algorithm. The path *p* that is currently being generated is shown in dark blue. The *i*-coverable frontier sub-path is indicated in yellow. The product of the out-degrees on this path (1 3 2 = 6) is smaller than *i*.

## 2 Background

### 2.1 Notation

A *sequence S* of length *n* is a tuple *S* ∈ Σ^*n*^ where Σ is a finite set Σ ={0,*…,σ–*1} called *alphabet*. The alphabet set for a DNA sequence can be defined as Σ_DNA_ = {A, C, G, T, N}, where N represents an unknown or ambiguous nucleotide. Since we primarily focus on DNA sequences in this study, the alphabet set is assumed to be the nucleotide alphabet denoted by Σ for simplicity throughout the article. The *i*th element of the sequence can be referred to as *s*_*i*_ and the sequence can be represented by concatenating all its elements *s*_0_*s*_1_ *… s*_*n* 1_. The *text string T* is a sequence terminated by a sentinel $ ∉ Σ. A *substring* of sequence *S* is indicated by *S*[*i… j*] = *s*_*i*_ *… s*_*j*_. The substring *S*[0 *… j*], and *S*[*j … n–*1] are called *prefix* and *suffix* of *S* and are denoted by *S*[*… j*] and *S*[*j …*], respectively. The term *k*-mer refers to any substring of length *k* in a string.

### 2.2 Sequence Graphs

Given an alphabet Σ, a tuple *G* = (*V, E, λ*) is a *sequence graph* over Σ; where *V* = {*v*_1_,*…, v* _|*V*|_ } is a set of *nodes, E* ⊆ *V × V* is a set of directed edges, and *λ* : *V*Σ^*^ is a function that maps each node in the graph to a *label* (see Figure 3a). We define 𝓁(*v*) := |*λ*(*v*)| as a short-hand for the label length of node *v*. We additionally assume that the graph is “deterministic”, in the sense that two outgoing edges starting at the same node are assumed to never target two nodes whose labels start with the same character. For a given node *v* ∈ *V*, the *out-degree* of *v* is the number of outward edges from *v*, denoted by *G*_*out*_(*v*), and *in-degree G*_*in*_(*v*) is the number of incoming edges. Any base *c* in the graph can be located by a tuple (*v, o*) where *v* ∈ *V* is the corresponding node containing *c* and *o* ∈ {0,*…,* 𝓁(*v*)–1} is the offset of *c* in *λ*(*v*). We call such a tuple *l* = (*v, o*) a *locus* in the graph.

**Figure 3.**
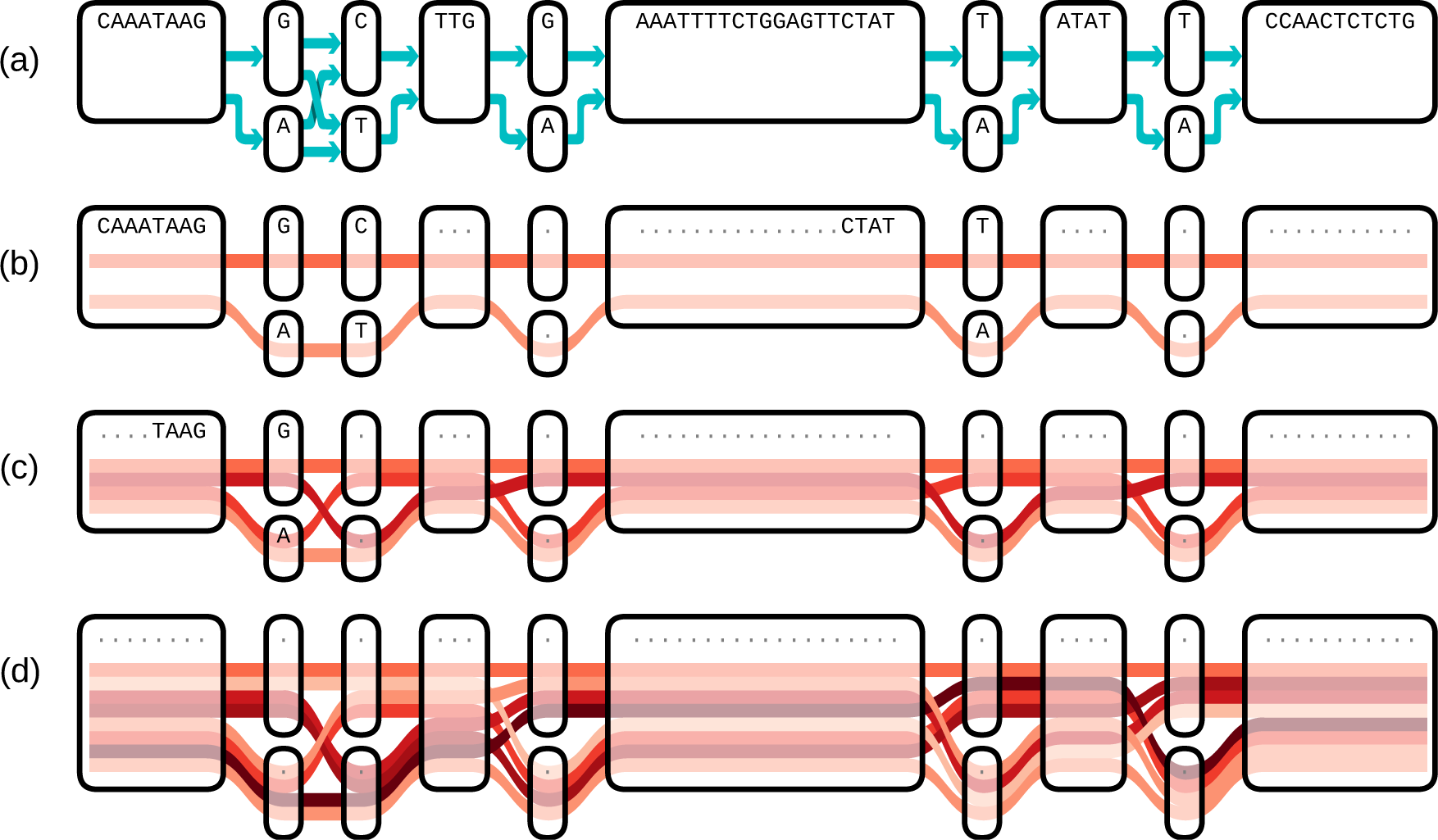
(a) Sequence graph with nodes displayed as boxes and edges indicated by blue arrows. The remaining three panels show the result of our path selection algorithm for different numbers of paths: (b) *N* = 2, (c) *N* = 4, and (d) *N* = 8. Selected paths are represented by red lines and *k*-covered loci for *k* = 10 are shown by dots.

In the literature, sequence graphs are sometimes defined such that each node implicitly represents a sequences and its reverse complement. For simplicity, we stick to the simpler definition here and consider the graph representing only the forward strand. However, it can be easily extended to bi-directed sequence graphs. Alternatively, this complication can be avoided by additionally querying the reverse complement seeds.

A path *P* in the graph is a sequence of nodes (*u, …, w*); where any two consecutive nodes in the path are connected by an edge in the graph. We define the sequence corresponding to the path *P* as the concatenation of its nodes: *λ*(*P*) = *λ*(*u*) *… λ*(*w*). A path in the graph that starts at offset *l* of the first node and ends at offset *r -* 1 of the last node is indicated by 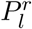.

The sequence graph of a species usually consists of multiple connected components corresponding to the multiple chromosomes. For each connected component *M*, we augment the graph with two additional nodes: a *head node h*_*M*_ and a *tail node t*_*M*_ with label *λ*(*h*_*M*_) = *λ*(*t*_*M*_) = ϵ, where ϵ denotes the empty string. We also add edges (*h, v*) for all *v* ∈ *V*_*M*_ which has zero in-degree and edges (*v, t*) for all *v* ∈ *V*_*M*_ that has zero out-degree. There are some special paths of interest: A *spanning path* of the component *M* is a path starting from head node *h*_*M*_ and ending in tail node *t*_*M*_. For any locus *l* = (*v, o*) in the graph *G* = (*V, E, λ*), a *k-path* is defined to be a path starting at *l* whose corresponding sequence length is *k*.

## 3 Methods

We consider a *set of reads R* ⊂ Σ^+^. First, a seed set *Q* is extracted from *R*:

### Algorithm 1 Path selection

**Require:** sequence graph *G* = (*V, E, λ*), path count *N*

1: **function** SELECTNEXTNODE(*p, P*)

2: *p′←p.*COVERABLEFRONTIERPATH(|*P*|)

3: *v ←p.*LASTNODE()

4: *cov*_min_ *←∞*

5: *v*_*c*_ *←*0

6: **for** *u* in *G.*ADJACENT(*v*) **do**

7: *cov ←P.*COVERAGE(*p′ u*)

8: **if** *cov*_min_ *> cov* **then**

9: *cov*_min_ *← cov*

10: *v*_*c*_ *← u*

11: **report** *v*_*c*_

12: **function** SELECTPATHS(*G, N*)

13: *P←* empty set

14: **while** *P* contains less than *N* paths **do**

15: *p ←G.*HEADNODE()

16: **while** *p* can be extended **do**

17: *c←* SELECTNEXTNODE(*p, P*)

18: *p.*APPEND(*c*)

19: **report** *P*

**Definition 1**

(Seed set). Given the set *R* of reads sequences, a length *k >* 0 and a distance *d >* 0. The *seed set Q*_*k,d*_(*R*) is defined as the set of all *k*-mers starting at positions *md* in the read sequences for any *m* ∈ ℕ_0_.

Note that for *d* = 1, the seed set *Q*_*k,d*_(*R*) is simply the set of all *k*-mers in reads *r* ∈ *R*, while for *d* = *k* it contains all *non-overlapping k*-mers. We now formalize the problem of seed finding as follows.

**Problem 2**

(Seed finding). Given a set *R* of read sequences, a sequence graph *G*, and parameters *k >* 0, *d >* 0. Find all occurrences of seeds *q* ∈ *Q*_*k,d*_(*R*) in paths in the graph *G*, where *Q*_*k,d*_(*R*) is the seed set according to Definition 1.

As discussed above, seed finding is a filtering strategy to limit the search space of sequence alignment algorithms. The choice of the seed length *k* controls the trade-off between specificity and sensitivity of this filter. Longer seeds increase specificity while reducing sensitivity. In this paper, we assume the value of *k* to be given as a parameter, which is usually chosen dependent on read length and error rate of the underlying sequencing technology.

### 3.1 Path Index

In the preprocessing phase, we create a *path index* of the genome graph (Figure 1, left/dark blue). The path index is essentially a compressed full-text index of a set of selected paths through the graph. Once constructed, it can be re-used for fast queries to find exact matches on these paths. The number of paths *N* is a tuning parameter of our indexing strategy; it can range from zero, which turns the path index off and seed finding happens purely in the traversal phase, to high numbers that lead to covering every *k*-path present in the graph. Constructing the path index proceeds in multiple steps:

#### Path Selection

The first step for constructing the path index is selecting a set of *N* paths. This step aims to cover as many *k*-paths in the graph as possible.

**Definition 3**

(Path Set Coverage). A path *p′* is *covered* by another path *p* if the node sequence of *p′* is a contiguous subsequence of the node sequence of *p*. We can generalize this to path sets: A set of paths *P covers* another path *p′* if and only if there is a path *p* ∈ *P* such that *p* covers *p′*. We define the *coverage* of path *p′* by set *P* as the number of paths in *P* that cover *p′*.

**Definition 4.**

A set of paths *P* in the graph *k-covers* a locus *l* = (*v, o*) if and only if, for all *k*-paths *p* starting at *l*, there is a path in *P* that covers *p*.

Based on Definition 4, *P* partitions the loci in the graph into two sets of *covered* and *uncovered* loci (for a given value of *k*). Every uncovered locus lowers the sensitivity of our path index. In order to reach full *p*sensitivity, all uncovered loci later need to be visited in the *graph traversal* phase, which we discuss below in Section 3.3. Consequently, maximizing the number of loci covered by the *N* selected paths minimizes the number of loci to be traversed. On the other hand, longer paths, ideally spanning paths, represent all covered *k*- paths in a more memory efficient way than shorter paths covering the same set of *k*-paths. So, the goal of path selection is finding a subset of *P ⊂ U* such that the number of loci covered by *P* would be maximized, where *U* is the set of all possible paths that start from the head node *h* and end at tail node *t*. More precisely, we seek to select *N* paths for each connected components of the graph (e.g. corresponding to the different chromosomes), resulting in a set *P* = {*p*_1_, *p*_2_,*…, p*_*Nm*_}, where *m* is the number components in the graph. For the sake of simplicity (and without loss of generality), we assume that the graph has only one component in the following.

We propose a heuristic greedy algorithm for selecting a set of paths that aims at covering as many loci as possible. This algorithm assumes that the input sequence graph is a *directed acyclic graph* (DAG). Although we do not pursue this further in this paper, the ideas we present could be extended to cyclic graphs, for instance by locally “unrolling” the graph into a DAG as done by VG (Garrison et al., 2018).

Our algorithm starts from an empty set *P*, and proceeds by incrementally adding paths to the set until it contains the desired number of paths. The basic idea is simple: To select an additional path, we walk the graph from the head to the tail node and greedily try to cover sub-paths that we have not covered before. That is, we want to extend a new path *p* such that it contains a little piece that is not yet covered by any other path selected thus far. To do this, we examine a local window around the present end of our new path *p* and refer to this window as the *frontier sub-path* (Figure 2). But how far should we look back, i.e. how many nodes should be included in this frontier sub-path? To determine this, we use a heuristic that reflects how many paths might possibly exist in this local neighborhood: We call a path *i-coverable*, if the product of the out-degrees of the nodes in that path is at most *i*. Assume that we have selected *i* paths so far and *p* is the (*i* + 1)-th path that we are presently selecting. We consider the shortest frontier sub-path of *p* that is *i*-coverable. Assume that *v* is the last node in *p*. To decide which node to append to *p* (i.e. choosing between *u* and *w* in Figure 2), we consider the paths selected so far and determine their coverage (see Definition 3) of the frontier sub-path extended by each node adjacent to *v*. We then choose the node with the lowest coverage, where ties are broken randomly.

Pseudocode for the path selection algorithm is given in Algorithm 1. We have visualized the outcome of this path selection algorithm for different numbers of paths in Figure 3 by using Sequence Tube Maps (https://github.com/vgteam/sequenceTubeMap). This greedy choice ultimately enumerates all paths in the graph for large values of *N*, while prioritizing them such that the first *m* paths aim to cover as many *k*-paths as possible in the graph. For simple graph topologies, such as the ones shown in Figure 3, this strategy maximizes the number of covered *k*-paths.

#### From Paths to Patches

The set of selected paths *P* can contain redundancies. That is, there can be sub-paths shared between multiple paths in *P*. This redundancy not only affects the memory footprint of our index, it also slows down the process of locating the occurrences in the graph. We therefore modify the selection procedure to generate *compressed* path sequences that avoid duplicate sub-paths. To this end, we define a parameter *T*, named *context length*. The idea is to produce the same paths as before but to “cut out” any redundant sub-paths while ensuring that all sub-paths of size *T* remain represented. Therefore, we add paths one-by-one and, before adding each path, determine which of its length-*T* sub-paths are not yet contained in any other path and refer to these sub-paths as *novel*. We remove all parts of the path that do not overlap with such a *novel* sub-path, and hence retain a set of disconnected *patches* of the path. This procedure ensures that all queries for strings of length at most *T* will remain unaffected. As a result, we usually obtain few genome-wide paths along with many smaller paths, the *patches*, cover variation sites.

#### Indexing

To build an index, we concatenate the set of these patches to form one sequence *S*_*P*_ wherein the patches *p*_*i*_ and *p*_*i*+1_ are separated by a sentinel $_*i*_ and *S*_*P*_ is terminated by $_*M* 1_, with $_*i*_ ∉ Σ_DNA_ for 0 *≤ i < M*. Then, we construct an FM index of *S*_*P*_ (Ferragina and Manzini, 2005), which we refer to as *path index*. The path index is accompanied by a two auxiliary data structures that allow us to later locate the nodes (and offsets within the nodes) for seed hits on these paths, which we will refer to as the LOCATE operation in Algorithm 2. This is facilitated by a data structure to support rank/select queries and a self-delimited integer vector encoded by Elias delta for storing the node IDs. The constructed path index can be used effectively to query any string shorter than the context length *T*. Note that smaller values of *T* usually lead to a smaller path index. In practice, we set *T* to the length of the seed hits we want to query. For the path index and the auxiliary data structures, we rely on the implementations available as part of the sdsl-lite library (Gog et al., 2014).

### 3.2 Chunk Index

One of the central ideas to enable full-sensitivity seed finding consists in processing the input read set *R* in chunks *R*_1_, *R*_2_,*…, R*_*C*_ ⊂ *R* and finding all seeds within a chunk simultaneously. To achieve this, we build an index over all seeds we want to query for the present read chunk *R*_*c*_, that is, we index the seed set *Q*_*k,d*_(*R*_*c*_) as introduced in Definition 1 (Figure 1 right/red). The underlying index structure could be a suffix tree, enhanced suffix array, or an FM index as long as top-down traversal operations are supported. After some preliminary experimentation, we decided to employ suffix trees constructed in a write-only top-down (WOTD) manner (Giegerich and Kurtz, 1995), since we observed best performance in practice and can tolerate the larger memory footprint compared to an FM index. WOTD trees are *lazy* suffix trees that are constructed during traversal and only evaluate parts of the tree that are actually traversed. Our key motivation for proceeding in chunks—rather than indexing the full read set—is rooted in the idea that the index over a chunk can fit in the processor cache (e.g. in L3 cache) and hence can answer queries swiftly in practice.

### 3.3 Traversing Graph, Path Index, and Chunk Index

Given a set of reads *R*, a seed length *k* and a seed distance *d*, our goal is to find all seed hits in a sequence graph *G* = (*V, E, λ*), that is, to solve Problem 2. For this purpose, we propose a novel strategy that proceeds in two phases. First, we perform a simultaneous traversal of path index and chunk index, yielding all seed hits represented in the selected paths. Second, we perform a simultaneous traversal of all uncovered graph loci and the chunk index, yielded seed hits missed by the path index, typically in variant-dense regions of the graph.

Even though represented by quite different data structures, sequence graph, path index and chunk index support a common set of abstract traversal operations. In the following, we describe our method in terms of such abstract operations and refer the reader to excellent text books on the details of these data structures (Ohlebusch, 2013; Mäkinen et al., 2015) as well as to mature implementations in libraries such as Seqan (Döring et al., 2008; Reinert et al., 2017) and SDSL (Gog et al., 2014).

More concretely, all three data structures (graph, path index, and chunk index) can be traversed using the following three operations:

- INITTRAVERSAL returns an initial *traversal location* 𝓁,
- ADVANCE(𝓁, *σ*) starts from traversal location 𝓁, consumes character *σ*, outputs the resulting location 𝓁′ or ∅ in case reading *σ* from location 𝓁 is not possible,
- EXTENSIONS(𝓁) returns the set of possible characters *σ* for which Advance(𝓁, *σ*) ≠∅.

In case of the suffix tree for the chunk index, a *traversal location* is described by a suffix tree node and an offset inside the node label, INITTRAVERSAL returns the root node and ADVANCE walks down the tree along the corresponding labels. For an FM index (which we use as path index), a *traversal location* is usually characterized by an interval in the Burrows-Wheeler transform, but just like for a suffix tree, traversal can be implemented such that ADVANCE returns a non-empty location as long as the spelled string is a substring of the indexed text. For the graph, a traversal can start from any locus and INITTRAVERSAL therefore needs to be supplied with a graph locus (which consists of node *v* and offset *o*, as described above). ADVANCE then walks along the graph in accordance with the node labels.

#### Algorithm 2 Finding seed hits *on* paths by simultaneous traversal of path index and chunk index

**Require:** chunk index CI, path index PI, length *k*

1: **function** FINDSEEDSONPATHS(CI, PI, *k*)

2: 𝓁_CI_ ←CI.INITTRAVERSAL()

3: 𝓁_PI_ ←PI.INITTRAVERSAL()

4: *states ←*empty queue

5: *states.*PUSH((𝓁_CI_, 𝓁_PI_, 0))

6: **while** *states* is not empty **do**

7: (𝓁_CI_, 𝓁_PI_, *k′*) ←*states.*Pop

8: **if** *k′*+1 *< k* **then**

9: Σ_ext_ ←CI.EXTENSIONS(𝓁_CI_) *∩* PI.EXTENSIONS(𝓁_PI_)

10: **for** *σ* in Σ_ext_ **do**

11: 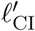 ← CI.ADVANCE(𝓁_CI_, *σ*)

12: 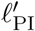←PI.ADVANCE(𝓁_PI_, *σ*)

13: *states.*PUSH 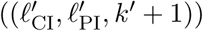

14: **else**

15: **report** PI.LOCATE(𝓁_PI_) *×* CI.LOCATE(𝓁_CI_)

#### Phase 1: Finding Seeds on Paths

To find seeds represented on the selected paths, we can simply query the path index. While, in principle, one could query each seed separately, we prefer to query all seeds in a chunk of reads at once through the simultaneous traversal of chunk index and path index (Algorithm 2 and Figure 1, Box 1). In this way, we benefit from the same chunk index that is also used in Phase 2 described below. Algorithm 2 assumes that the path index supports the additional operation LOCATE, which translates a traversal location in the path index into a set of corresponding locations in the graph (see Section 3.1).

#### Phase 2: Finding Seeds off Paths

Since the path index can occasionally miss *k*-paths, we handle variant-dense parts of the graph in a second phase. During the selection of paths, we keep track of loci that are not covered by paths and store the set *L* of these uncovered loci. For each chunk, we examine all these uncovered loci and, starting from these loci, simultaneously traverse the graph and the chunk index (Algorithm 3 and Figure 1, Box 2). In this way, the seeds that are contained in the chunk of reads guide the traversal of the graph. This allows us to avoid enumerating all *k*-paths at the uncovered loci, which would be infeasible. For traversing an uncovered graph locus, the number of ADVANCE operations is bounded by the size of the chunk sequence, i.e. *C · m*; where *C* is the number of reads in the chunk, and *m* is the average reads length. Thus, the total time complexity of finding seeds off paths is *O*(|*L*|*· C · m ·* Σ), where *L* is the set of uncovered loci.

##### Algorithm 3 Finding seed hits *off* paths by simultaneous traversal of graph and chunk index

**Require:** chunk index CI, graph *G* = (*V, E, λ*), set of uncovered locations *L*, length *k*

1: **function** FINDSEEDSOFFPATHS(CI, *G, L k*)

2: **for** 𝓁_*G*_ in *L* **do**

3: 𝓁←𝓁_*G*_

4: 𝓁_CI_*←* CI.INITTRAVERSAL()

5: *states ←* empty queue

6: *states.*Push((𝓁_CI_, 𝓁_*G*_, 0))

7: **while** *states* is not empty **do**

8: (𝓁_CI_, 𝓁_*G*_, *k′*) *←states.*Pop

9: **if** *k′* +1 *< k* **then**

10: Σ_ext_ *←* CI.EXTENSIONS(𝓁_CI_) *∩* G.EXTENSIONS(𝓁_G_)

11: **for** *σ* in Σ_ext_ **do**

12: 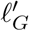 *← G.*ADVANCE(𝓁_*G*_, *σ*)

13: 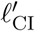 *←*CI.ADVANCE(𝓁_CI_, *σ*)

14: *states.*PUSH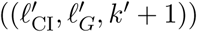

15: **else**

16: **report** {𝓁}*×* CI.LOCATE(𝓁_CI_)

## 4 Experimental Results

### 4.1 Implementation

PSI has been implemented in C++. It gets the reads set in `FASTQ` and the graph in `vg` format as inputs, and finds occurrences of all seeds with given length *k* and distance *d*. The output is provided in `GAM` format which represents seed alignments to the graph. Both `vg` and `GAM` are file formats introduced by the VG toolkit to represent sequence graph and sequence alignment, respectively (Garrison et al., 2018). In order to maintain interoperability between tools in this domain, we reuse these file formats. For internal usage, the graph is represented by `xg`, the succinct graph data structure of VG, which allows to access node sequences and connectivities efficiently. We use the `sdsl-lite` library (Gog et al., 2014) for succinct and compressed data structures: bit vector with efficient `rank` and `select` operations, compressed integer vector using Elias delta coding, and FM index. The WOTD-tree we use is provided by the SeqAn2 library (Reinert et al., 2017).

All running times are measured on a system with a 3 GHz Intel Xeon E7-8857 processor running Debian 9.4 with Linux kernel 4.9.91. We used `libvg` version 1.7.0 and `sdsl-lite` version 2.1.1 for the benchmarks. Seed finding is done using a single thread.

### 4.2 Data Sets

We benchmark our algorithm using both synthetic and real graphs. In both cases, we start from a linear reference genome and a set of small variants and use VG version 1.7.0 (Garrison et al., 2018) to construct a corresponding graph (using `vg construct` command). This process results in a DAG with one bubble for each implanted variant.

#### Simulated Graphs

To systematically explore parameter settings and to benchmark the performance across a wide range of graphs with different complexities, we created a simulated data set. This data set is constructed from the complete genome of *N. Deltocephalinicola*, a bacterial species with a short genome of around 112 kbp (Bennett et al., 2016). Starting from this linear genome, single nucleotide variants (SNVs) are implanted uniformly at random throughout the genome with three different mutation rates (0.01, 0.1, 0.3) to obtain three graphs ranging from moderate variant density (0.01) to an extreme variant density (0.3).

#### 1000 Genomes Graphs

The real data set consists of graphs constructed from the autosomes of the human reference genome (hs37d5) and small variants reported by Phase 3 of the 1000 Genome Project (Auton et al., 2015). We created two versions of this graph, one constructed from all small variants, and a second one that only includes variants with an allele frequency above 1%. Statistics for the resulting graphs are reported in Table 1.

**Table 1.**
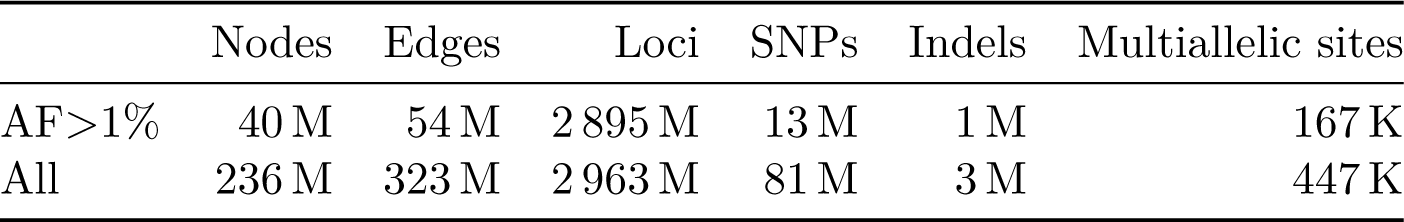
Human genome variation graph statistics.

#### Read Simulation

To benchmark our seed finding method, we simulated one million reads of length 150 bp with error rate 1% for each graph. The reads were simulated from random haplotypes, created by a random walk through the respective graph. The number of haplotypes used for read simulation corresponds to the ploidy of underlying genome: one haplotype for the simulated graph (bacteria) and two haplotypes for the 1000 Genomes graphs (human). During seed finding, we query all non-overlapping seeds of length 30 bp.

### 4.3 Performance on Simulated Graphs

We used the controlled environments provided by the simulated graphs to comprehensively explore the properties of our path index when confronted with graphs of varying variant densities.

#### Index size

First, we examined the influence of the number of paths *N* on the path index size. When turning off the path compression, that is, indexing all paths in full without removing redundant parts, then the index size increases linearly in the number of paths (Figure 4, red curves). As shown by the cyan curves, the compression/patching considerably decreases the size of the index, particularly for simple graphs. In complex graphs, paths share fewer identical substrings that can be dropped. Another factor that affects the compression rate is the *context length T*. Higher value of *T* result in longer patches which leads to a bigger index. Figure 4 includes results for two context lengths 32 and 64. Recall that the *context length* imposes an upper limit for query pattern length, where 32 constitutes a typical value for seed finding in Illumina short reads.

**Figure 4.**
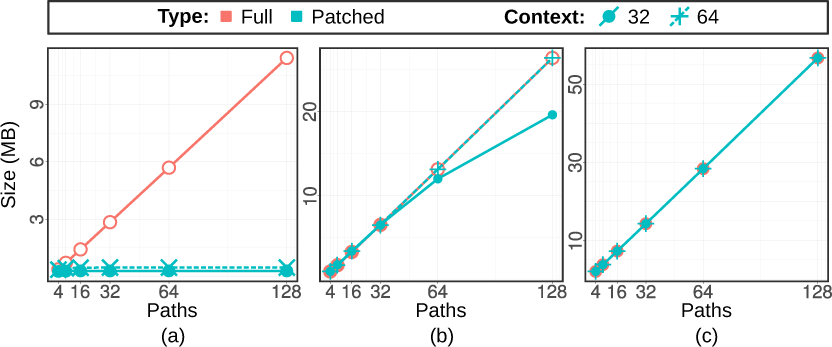
Compressed (patched) path index size in MB vs. different number of paths with context length 30 compared to uncompressed (full) one for simulated data set with mutation rates: (a) 0.01, (b) 0.1, and (c) 0.3.

#### Indexing time

Figure 5 shows the time spent on different phases of creating the path index, namely on path selection (“pick”) on creating the FM index (“index”) and on writing the index to disk (“save”). As Figure 5 shows, the times spent on the indexing phase are dominated by the path selection phase, while the time for saving is negligible. The growth of the runtime of the path selection is slightly super-linear in practice.

**Figure 5.**
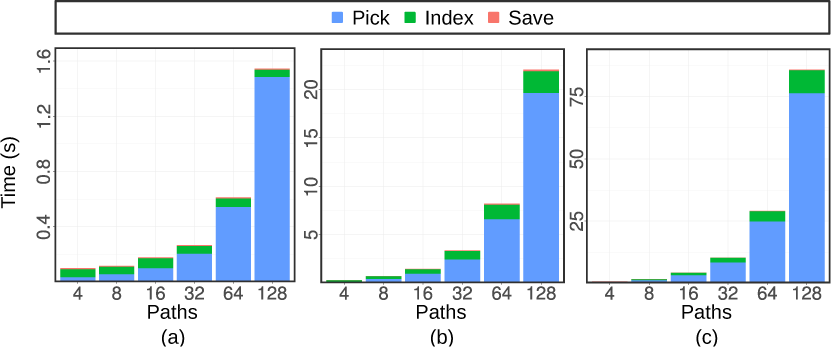
Time spent on different phases of path indexing for simulated data set with mutation rates: (a) 0.01, (b) 0.1, and (c) 0.3.

**Figure 6.**
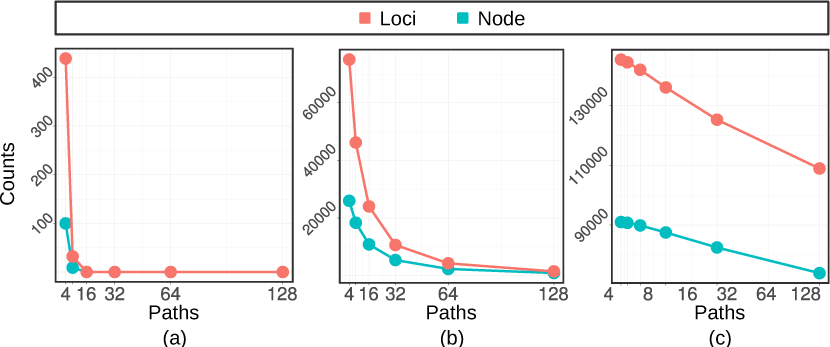
Number of uncovered loci/nodes by indexes with different number of paths for simulated data set with mutation rates: (a) 0.01, (b) 0.1, and (c) 0.3.

#### Path coverage

The efficiency of the path selection algorithm in terms of covering *k*-mers (for *k* = 30, referred to as “loci”) and graph nodes is plotted in Figure 6. The number of uncovered loci is shown for different sizes of the path set *P*. These curves show a behavior that is consistent with the distribution of the number of SNPs covered by each *k*-mers (Figure 7). For the intermediate SNP density of 0.1 (middle), for example, we expect a 30-mer to cover 3 SNPs on average, which translates into 2^3^ = 8 paths needed to cover all “versions” of this 30-mer.

**Figure 7.**
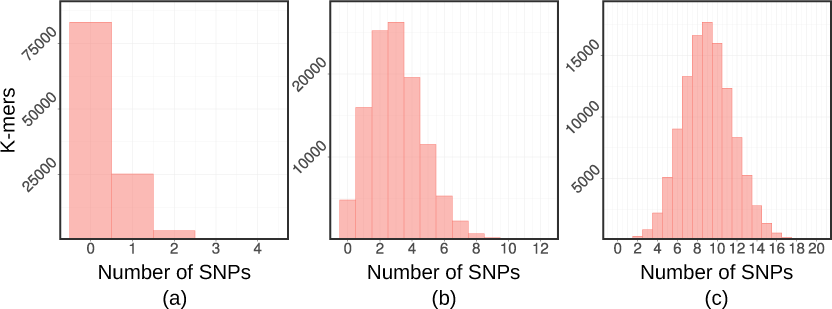
Histogram of SNPs frequency in 30-mers for simulated data set with mutation rates: (a) 0.01, (b) 0.1, and (c) 0.3.

#### Seed Finding

We now employ the hybrcid index using both the stages of querying the path index and traversing the graph to recover seeds that are missed by the path index. We measure the total runtime of both phases. To make the numbers comparable to the human data, where the same seed can sometimes occur many times, we divide the total runtime by the number of occurrences found to obtain the average runtime per seed query. Figure 8 shows the resulting seed query times for the three graphs as a function of number of selected paths and the chunk size. In line with our expectation, the query time decreases when adding more paths and when increasing the chunk size. For more variant-rich graphs, the queries become slower.

**Figure 8.**
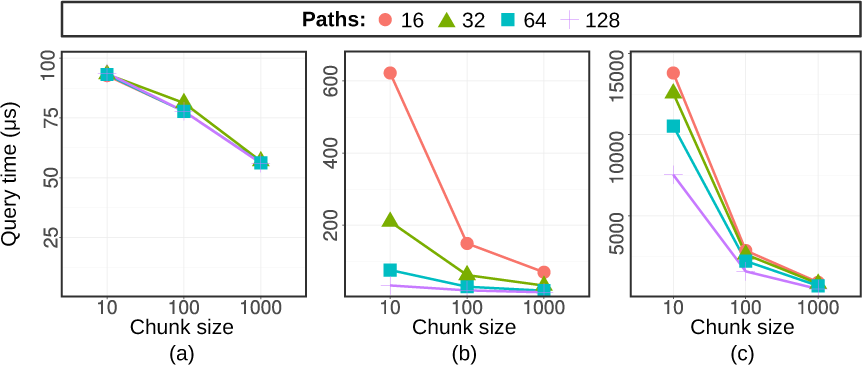
Average query time per *k*-mer occurrence for different numbers of paths and chunk sizes (given in the number of reads/chunk) on simulated data set with mutation rates: (a) 0.01, (b) 0.1, and (c) 0.3.

### 4.4 Performance on 1000 Genomes Graphs

Experiments on the large 1000 Genomes graphs reveal that the path index behaves similarly favorable as for the small simulated graphs. Figure 9 shows different measurements for the graph with all variants with allele frequency of 1% and above. We observe that our path compression (patching) routine is very effective in limiting the size of the index. Even when indexing patches corresponding to 256 paths through the full human genome, we observe path index sizes below 7GB (Figure 9a). While we see the same super-linear growth in runtime as for the simulated graphs, the construction of the path index is easily feasible, with less than 10 hours for 128 paths and less than 30 hours for 256 paths (Figure 9b). Again, the number of uncovered *k*-mers is quickly driven down by adding more paths (Figure 9c).

**Figure 9.**
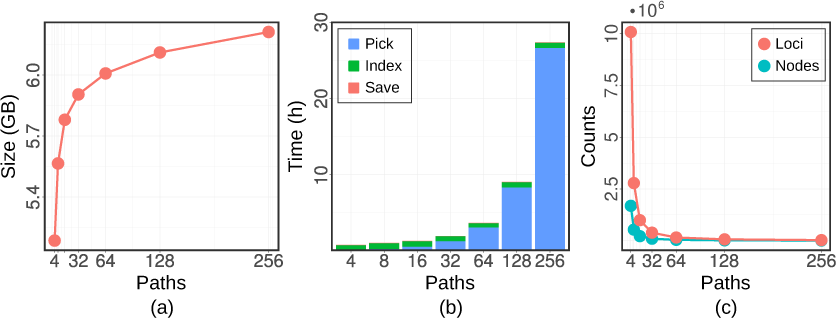
Human genome path index benchmark for different number of paths: (a) path index size, (b) indexing time, and (c) number of uncovered loci/nodes

In Figure 10, we examine the dependency of the *k*-mer query performance on the chunk size, which reveals that finding seeds in chunks of 100 000 reads is most favorable. The performance becomes worse when using even larger chunks, which we attribute to cache effects.

**Figure 10.**
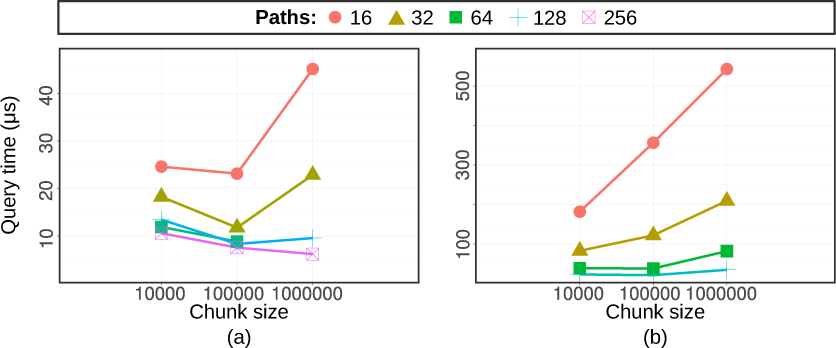
Average query time per *k*-mer for different number of paths and chunk sizes (given in the number of reads/chunk) on the human genome data set: (a) allele frequency above 1%, (b) all variants.

Finally, we compare the performance to GCSA2, a state-of-the-art method for indexing graphs developed by Sirén (2017) and used in VG (Garrison et al., 2018). The results are displayed in Table 2. For the graph with variants of AF *>* 1%, we obtain an index less than half the size (6.3 GB) of that produced by GCSA2 (15 GB), while we only need slightly longer to construct it (28h vs. 22h). Our path index (“PSI/Path-only”) covers more *k*-mers (99.24% vs. 99.09%) and allows for faster queries, 4.8*μs* per occurrence where GCSA2 needs 6.28*μs* per occurrence—a speedup of 30.8%. When additionally using the graph traversal (“PSI/Hybrid”) to rescue the uncovered *k*-mers, our query time is virtually the same as GCSA2 while reaching full sensitivity, which is not feasible with GSCA2.

**Table 2.** Seed finding performance on the 1000 Genomes graphs.

|  |  | Index size | Indexing time | Covered k-mers | Query time |
| --- | --- | --- | --- | --- | --- |
| AF > 1% | PSI/Path-only | 6.3 GB | 28 h | 6 721 M (99.24%) | $4.82 \mu s / occ$ |
| | PSI/Hybrid | 6.3 GB | 28 h | 6 773 M (100%) | $6.20 \mu s / occ$ |
| | GCSA2 order-128 | 15 GB | 22 h* | 6 712 M (99.09%) | $6.28 \mu s / occ$ |
| All | PSI/Path-only | 64 GB | 56 h | | $8.12 \mu s / occ$ |
| | PSI/Hybrid | 64 GB | 56 h | 89 207 M (100%) | $21.05 \mu s / occ$ |
| | GCSA2 order-128 | 34 GB | 30 h* | 13 863 M (15.54%) | $20.03 \mu s / occ$ |

The graph with all variants contains drastically more *k*-mers (8.9 10^10^) than the graph with variants of AF *>* 1% (6.7 10^9^). In this setting, the pruning steps required to build the GCSA2 index (which we run as described in the GCSA2 documentation) lead to a drastic loss in the number of indexed *k*-mers: the GCSA2 index only captures 15.54% of all *k*-mers in the graph. Even though the lost *k*-mers are concentrated in the complex regions of the graph, we argue that making such regions accessible is one important objectives of switching from linear reference genomes to graphs in the first place. Using PSI/Hybrid, we reach full sensitivity for this graph with a comparable query time (21.05*μs* for PSI/Hybrid and 20.03*μs* for GCSA2).

## 5 Discussion

We have introduced an approach to index sequence graphs that scales to human genomes while delivering full sensitivity. Our path selection procedure coupled with an FM index results in a competitive index structure, even when used in isolation without the graph traversal phase. By traversing the graph and the chunk index simultaneously, we take advantage of the fact that the set of *k*-mers in the reads is more restricted than the one represented in the graph. In other words, we let the reads guide the traversal of the graph and, in this way, circumvent the combinatorial explosion of *k*-mers in the graph. For the first time, this techniques enables *scalable full-sensitivity seed finding in variation graphs*.

Here, we focused on introducing a new algorithmic technique for finding seeds in variation graphs. Our results show that full-sensitivity seed finding is indeed possible in polynomial time and that it can be done efficiently in practice. We plan to use this method to build a full read mapper by combining it with our recent algorithm for bit-parallel sequence-to-graph alignment (Rautiainen et al., 2019).

Recently, Sirén et al. (2018) have proposed to augment sequence graphs with paths that represent haplotypes found in a population, to then restrict the indexing to those haplotypes. This idea could naturally be combined with our method by replacing the path selection step accordingly, which we plan to explore in future research. Beyond that, Pritt et al. (2018) have argued that it might be beneficial to restrict the set of variants used for graph construction to a well-selected subset for two reasons: to avoid introducing unnecessary ambiguity and to simplify indexing. By providing a full-sensitivity index, we have removed the necessity for the latter, creating the opportunity for comprehensive evaluations on the trade-off between added ambiguity and reduced read mapping bias.

## 6 Acknowledgement

We thank Mikko Rautiainen for pointing out the polynomial time complexity of finding seeds off paths.

